# The status of the microbial census: an update

**DOI:** 10.1101/038646

**Authors:** Patrick D. Schloss, Rene Girard, Thomas Martin, Joshua Edwards, J. Cameron Thrash

## Abstract

A census is typically carried out for people at a national level; however, microbial ecologists have implemented a molecular census of bacteria and archaea by sequencing their 16S rRNA genes. We assessed how well the microbial census of full-length 16S rRNA gene sequences is proceeding in the context of recent advances in high throughput sequencing technologies. Among the 1,411,234 and 53,546 full-length bacterial and archaeal sequences sequences, 94.5% and 95.1% of the bacterial and archeaeal sequences, respectively, belonged to operational taxonomic units (OTUs) that have been observed more than once. Although these metrics suggest that the census is approaching completion, 29.2% of the bacterial and 38.5% of the archaeal OTUs have been observed more than once. Thus, there is still considerable microbial diversity to be explored. Unfortunately, the rate of new full-length sequences has been declining and new sequences are primarily being deposited by a small number of studies. Furthermore, sequences from soil and aquatic environments, which are known to be rich in bacterial diversity, only represent 7.8 and 16.5% of the census while sequences associated with zoonotic environments represent 55.0% of the census. Continued use of traditional approaches and new technologies such as single cell genomics and short read assembly are likely to improve our ability to sample rare OTUs if it is possible to overcome this sampling bias. The success of ongoing efforts to use short read sequencing to characterize microbial communities requires that researchers strive to expand the depth and breadth of the microbial census.

**Importance:** The biodiversity contained within the bacterial and archaeal domains dwarfs that of the eukaryotes and the services these organisms provide to the biosphere are critical. Surprisingly, we have done a relatively poor job of keeping track of the ongoing effort to characterize the biodiversity as represented in full-length 16S rRNA genes. By understanding how this census is proceeding, it is possible to suggest the best allocation of resources for advancing the census. We found that the ongoing effort has done an excellent job of sampling the most abundant organisms, but struggles to sample the more rare organisms. Through the use of new sequencing technologies we should be able to obtain full-length sequences from these rare organisms. Furthermore, we suggest that by allocating more resources to sampling environments known to have the greatest biodiversity we will be able to make significant advances in our characterization of microbial diversity.

## Introduction

The effort to quantify the number of different organisms in a system remains fundamental to understanding ecology (1, 2). At the scale of microorganisms, small physical sizes, morphological ambiguity, and highly variable population sizes complicate this process. Furthermore, creating standards for delimiting what makes one microbe “different” from another has been contentious (3, 4). In spite of these challenges, we continue to peel back the curtain on the microbial world with the aid of more and more informative, if still limited, technologies like cultivation, 16S rRNA gene surveys, single cell technologies, and metagenomics.

Generating a comprehensive understanding of any system with a single gene may seem a fool’s errand, yet we have learned a considerable amount regarding the diversity, dynamics, and natural history of microorganisms using the venerable 16S rRNA gene. In 1983, the full-length 16S rRNA gene sequence of *Escherichia coli* (accession J01695) was deposited into NCBI’s GenBank making it the first of what is now more than 10 million 16S rRNA gene sequences to be deposited into the database (5). 16S rRNA gene accessions represent nearly one-third of all sequences deposited in GenBank, making it the best-represented gene. As Sanger sequencing has given way to so-called “next generation sequencing” technologies, hundreds of millions of 16S rRNA gene sequences have been deposited into the NCBI’s Sequence Read Archive. The expansion in sequencing throughput and increased access to sequencing technology has allowed for more environments to be sequenced at a deeper coverage, resulting in the identification of novel taxa. The ability to obtain sequence data from microorganisms without cultivation has radically altered our perspective of their role in nearly every environment from deep ocean sediment cores (e.g. accession AY436526) to the International Space Station (e.g. accession DQ497748).

Previously, Schloss and Handelsman (6) assigned the 56,215 partial 16S rRNA gene sequences that were available in the Ribosomal Database Project to operational taxonomic units (OTUs) and concluded that the sampling methods of the time were insufficient to identify the previously estimated 10^7^ to 10^9^ different species (7, 8). That census called for a broader and deeper characterization of all environments. Refreshingly, this challenge was largely met. There have been major investments in studying the Earth’s microbiome using 16S rRNA gene sequencing through initiatives such as the Human Microbiome Project (9), the Earth Microbiome Project (10), and the International Census of Marine Microorganisms (11). But most importantly, the original census was performed on the cusp of radical developments in sequencing technologies. That advancement has moved the generation of sequencing throughput from large sequencing centers to individual investigators and leveraged their diverse interests to expand the representation of organisms and environments represented in public databases.

It is disconcerting that the increase in sequencing volume has come at the cost of sequence length. The commonly used MiSeq-based sequencing platform from Illumina is extensively used to sequence the approximately 250 bp V4 hypervariable region of the 16S rRNA gene; other schemes have used different parts of the gene that are generally shorter than 500 bp. The number of OTUs that are sampled when using different regions within the 16S rRNA gene can vary considerably and the genetic diversity within these regions typically has only a modest correlation the genetic diversity of the full-length sequence (12, 13). Thus, it remains unclear to what degree richness estimates from short read technologies over or underestimate the numbers from full-length sequences. Furthermore, we likely lack the references necessary to adequately classify the novel biodiversity we are sampling when we generate 100-times the sequence data from a community than we did using full-length sequencing.

Here we update the status of the microbial census with full-length 16S rRNA gene sequences. In the 13 years since the collection of data for Schloss and Handelsman’s initial census, the number of full-length sequences has grown exponentially, despite the overwhelming contemporary focus by most researchers on short-read technologies. This update to the census allows us to evaluate the relative sampling thoroughness for different environments and clades and make an argument for the continued need to collection full-length sequence data from many systems that have a long history of study. As researchers consider coalescing into a Unified Microbiome Initiative (14), it will be important to balance the need for mechanism-based studies with the need to generate full-length reference sequences from a diversity of environments.

## Results and Discussion

### The status of the bacterial and archaeal census

To assess the field’s progress in characterizing the biodiversity of bacteria and archaea, we assigned each 16S rRNA gene sequence to OTUs using distance thresholds that varied between 0 and 20%. Although it is not possible to link a specific taxonomic level (e.g. species, genus, family, etc.) to a specific distance threshold, we selected distances of 0, 3, 5, 10, and 20% because they are widely regarded as representing the range of genetic diversity of the 16S rRNA gene within each domain. By rarefaction, it was clear that the ongoing sampling efforts have started to saturate the number of current OTUs. After sampling 1,411,234 near full-length bacterial 16S rRNA gene sequences we have identified 217,645, 108,950, 66,819, 15,743, and 3,731 OTUs at the respective thresholds (Figure 1A, Table 1). Using only the OTUs generated using a 3% threshold, we calculated a 94.5% Good’s coverage (percent of sequences belonging to OTUs that have been observed more than once), but only 29.2% OTU coverage (percent of the OTUs that have been observed more than once). Paralleling the bacterial results, after sampling 53,546 archaeal 16S rRNA gene sequences we have identified 11,040, 4,252, 2,364, 812, and 110 OTUs (Figure 1B, Table 1). Using only the OTUs generated with a 3% threshold, we calculated a 95.1% Good’s coverage, but only 38.5% OTU coverage. These results indicate that regardless of the domain, continued sampling with the current strategies for generating full-length sequences will largely reveal OTUs that have already been observed, even though a large fraction of OTUs have only been sampled once. Considering more than 70.8% of the OTUs have only been observed once, it is likely that an even larger number of OTUs have yet to be sampled for both domains.

**Figure 1.**
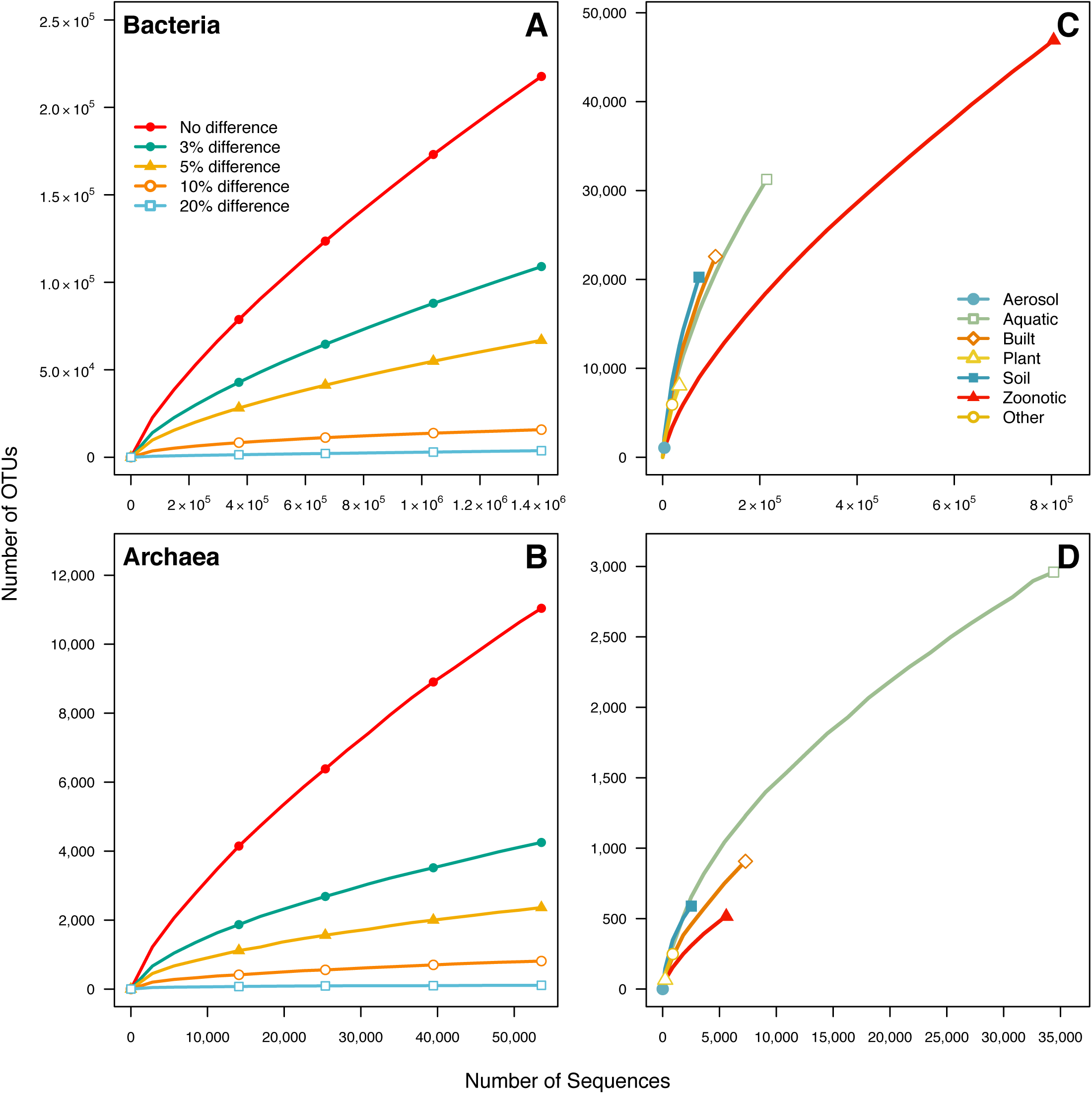
Number of OTUs sampled among bacterial and archaeal 16S rRNA gene sequences for different OTU definitions and level of sequencing effort. Rarefaction curves for different OTU definitions of Bacteria (A) and Archaea (B). Rarefaction curves for the coarse environments in Table 1 for Bacteria (C) and Archaea (D). The number of bacterial and archaeal OTUs observed among the longest sequences in the SILVA database continues to grow at a rate too slow to ever reach estimates of 10^6^ to 10^11^ bacterial species.

**Table 1.** Status of microbial census by habitat classifications and domain. The isolation_source field from the SILVA reference database was manually curated to assign bacterial and archaeal sequences coarse and fine scale habitat classifications. We calculated the number of sequences and OTUs observed and the percent coverage on a sequence or OTU basis for each classification and domain. Descriptions of each category are provided in Table S1.

| Coarse | Fine | Bacteria |  |  |  | Archaea |  |  |  |
| --- | --- | --- | --- | --- | --- | --- | --- | --- | --- |
|  |  | Sequences (N) | OTUs (N) | % Seq. Coverage | % OTU Coverage | Sequences (N) | OTUs (N) | % Seq. Coverage | % OTU Coverage |
| Aerosol |  | 3,472 | 1,068 | 79.5 | 33.2 | 2 | 1 | 100.0 | 100.0 |
| Aquatic | Brackish | 1,094 | 646 | 54.6 | 23.1 | 1,368 | 314 | 87.4 | 44.9 |
|  | Brackish sediment | 390 | 243 | 54.4 | 26.7 | 525 | 208 | 76.8 | 41.3 |
|  | Freshwater | 21,647 | 6,689 | 80.8 | 37.7 | 1,540 | 439 | 84.7 | 46.5 |
|  | Freshwater sediment | 6,733 | 3,549 | 63.0 | 29.8 | 1,324 | 488 | 79.3 | 43.9 |
|  | Marine | 134,727 | 14,287 | 94.3 | 46.7 | 10,983 | 830 | 95.8 | 44.5 |
|  | Marine sediment | 27,801 | 9,567 | 79.6 | 40.8 | 14,049 | 1,507 | 95.0 | 53.7 |
|  | Hydrothermal vent | 10,860 | 4,216 | 75.4 | 36.5 | 3,797 | 734 | 90.4 | 50.3 |
|  | Ice | 2,073 | 936 | 71.2 | 36.1 | 42 | 5 | 95.2 | 60.0 |
|  | Other | 8,760 | 3,802 | 71.7 | 34.8 | 772 | 313 | 80.7 | 52.4 |
| Built | Digesters | 33,152 | 8,949 | 82.9 | 36.8 | 4,764 | 483 | 93.6 | 36.4 |
|  | Food-associated | 11,813 | 1,632 | 92.0 | 41.9 | 117 | 40 | 80.3 | 42.5 |
|  | Industrial/mining | 16,582 | 6,099 | 76.6 | 36.3 | 1,245 | 336 | 84.4 | 42.3 |
|  | Pollution associated | 38,696 | 10,602 | 84.1 | 41.9 | 716 | 249 | 79.2 | 40.2 |
|  | Other | 8,556 | 2,730 | 79.1 | 34.7 | 444 | 111 | 90.8 | 63.1 |
| Plant associated | Root | 19,695 | 5,052 | 84.3 | 38.7 | 200 | 61 | 85.5 | 52.5 |
|  | Surface | 4,892 | 1,385 | 82.7 | 38.8 | 0 | 0 | NA | NA |
|  | Other | 9,753 | 3,217 | 78.8 | 35.8 | 22 | 7 | 90.9 | 71.4 |
| Soil | Agriculture | 10,051 | 4,017 | 73.6 | 34.0 | 146 | 56 | 80.8 | 50.0 |
|  | Desert | 3,042 | 1,280 | 73.7 | 37.5 | 245 | 79 | 77.6 | 30.4 |
|  | Permafrost | 1,922 | 870 | 73.0 | 40.3 | 39 | 20 | 64.1 | 30.0 |
|  | Other | 59,855 | 17,166 | 82.9 | 40.4 | 2,087 | 516 | 89.1 | 55.8 |
| Zoological | Vertebrate | 773,045 | 42,497 | 96.1 | 29.5 | 5,389 | 454 | 95.1 | 41.6 |
|  | Arthropod | 13,209 | 3,688 | 81.8 | 34.7 | 87 | 52 | 58.6 | 30.8 |
|  | Other invertebrate | 7,476 | 2,626 | 78.0 | 37.3 | 67 | 30 | 73.1 | 40.0 |
|  | Other | 10,855 | 1,754 | 89.2 | 33.4 | 54 | 17 | 87.0 | 58.8 |
| Other |  | 19,414 | 5,930 | 81.6 | 39.9 | 882 | 249 | 84.2 | 44.2 |
| No source data |  | 151,669 | 14,144 | 94.9 | 45.6 | 2,565 | 559 | 88.6 | 47.6 |
| Total |  | 1,411,234 | 108,950 | 94.5 | 29.2 | 53,546 | 4,252 | 95.1 | 38.5 |

### Sequencing efforts are a source of bias in the census

One explanation for the large number of OTUs that have only been observed once is that with the the broad adoption of sequencing platforms that generate short sequence reads, the rate of full-length sequence generation has declined. In fact, since 2009 the number of new bacterial sequences generated has slowed to an average of 189,960 sequences per year (Figure 2A). Although this is still an impressive number of sequences, since 2007 the number of new bacterial OTUs has plateaued at an average of 11,184 new OTUs per year (Figure 2B). Given the expense of generating full-length sequences using the Sanger sequencing technology and the transition to other platforms at that time, we expected that the large number of sequences were being deposited by a handful of large projects. Indeed, when we counted the number of submissions responsible for depositing 50% of the sequences, we found that with the exception of 2006 and 2013, eight or fewer studies were responsible for depositing the majority of the full-length sequences each year since 2005 (Figure 2C). Between 2009 and 2012, 908,190 total sequences were submitted and 6 submissions from 5 studies were responsible for depositing 550,274 (60.6% of all sequences). These studies generated sequences from the human gastrointestinal tract (15), human skin (16, 17), murine skin (18), and hypersaline microbial mats (19). The heavy zoonotic focus is reflected in the rarefaction curve for this category (Figure 1C). In contrast to recent years, between 1995 and 2006, an average of 39.3 studies were responsible for submitting more than half of the sequences each year. Although the recent deep surveys represent significant contributions to our knowledge of bacterial biogeography, their small number and lack of environmental diversity is indicative of the broader problems in advancing the bacterial census.

**Figure 2.**
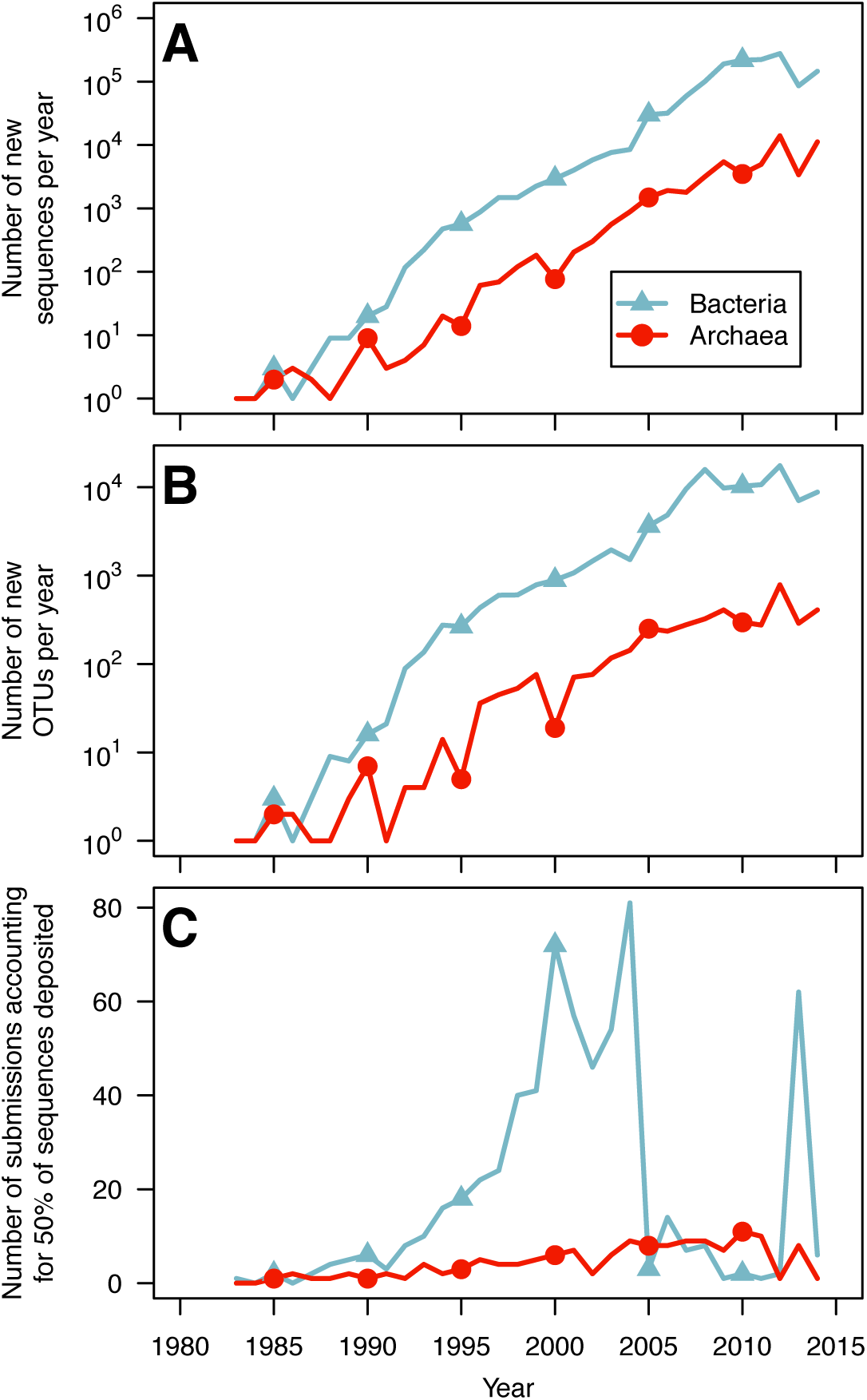
Progression of the microbial census since the first full-length 16S rRNA gene sequence was deposited into GenBank in 1983.* The number of bacterial and archaeal 16S rRNA gene sequences deposited (A) and the new OTUs they represent (B) has increased exponentially until the last several years when the rate of change has plateaued. For both bacterial and archaeal sequences, the number of studies that are responsible for depositing more than 50% of the sequences each year has been relatively small (C).

The depth of sequencing being done to advance the archaeal census has been 26-times less than that of the bacterial census (Table 1). The annual number of sequences submitted has largely paralleled that of the bacterial census with a plateau starting in 2009 and an average of 7,075 sequences each year since then. The number of new archaeal OTUs represented by these sequences began to slow in 2005 with an average of 355.5 new OTUs per year. With the exception of 2012 and 2014, the number of submissions responsible for more than 50% of the archaeal sequences submitted per year has varied between 2 and 11 submissions per year. The clear bias towards sequencing bacterial 16S rRNA genes has limited the ability to more fully characterize the biodiversity of the archaea, which is clearly reflected in the relatively meager sampling effort across habitats, compared to bacteria (Figure 1D),

### The ability to sample microbial life is taxonomically skewed

The Firmicutes, Proteobacteria, Actinobacteria, and Bacteroidetes represent 89.2% of the bacterial sequences and the Euryarchaeota and Thaumarchaeota 86.5% of the archaeal sequences. We sought to understand how the representation of individual phyla has changed relative to the state of the census in 2006. We used 2006 as a reference point for calibrating the dynamics of the bacterial and archaeal censuses since that was the year that the first highly parallelized 16S rRNA gene sequence dataset was published (20). Based on the representation of sequences within the SILVA database, in 2006 there were 61 bacterial and 18 phyla. Since then there have been 4 new bacterial (CKC4, OC31, S2R-29, and SBYG-2791) and 2 new archaeal candidate phyla (Ancient Archaeal Group and TVG8AR30). Relative to the overall sequencing trends before and after 2006, several phyla stand out for being over and underrepresented in sequence submissions (Figure 3). Among the bacterial phyla with at least 1,000 sequences, Atribacteria and Kazan-3B-09 were sequenced 4-fold more often while Deinococcus-Thermus and Tenericutes were sequenced 2-fold less often than would have been expected since 2006. Among the archaeal phyla with at least 1,000 sequences, the Thaumarchaeota were sequenced 2.0-fold more often and the Crenarchaeota were sequenced 6.7-fold less often than expected. Together, these results demonstrate a change in the phylum-level lineages represented in the census from before and after 2006 and encouragingly, show that some underrepresented phyla are becoming better sampled.

**Figure 3.**
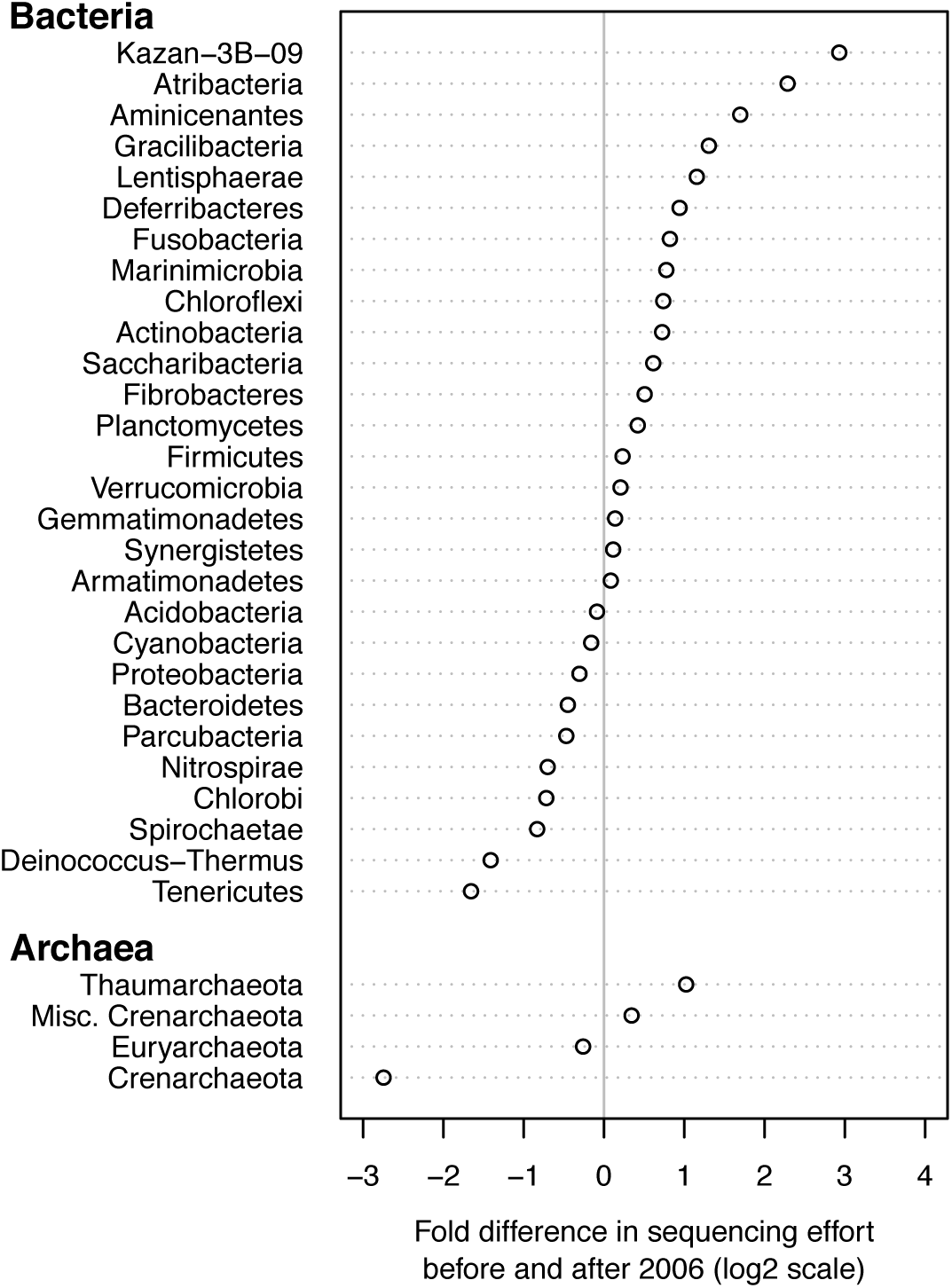
Relative rate of sequence deposition for each bacterial and archaeal phylum before and after 2006 relative to the sequencing of all bacteria. The figure shows the relative rates for those phyla with at least 1,000 sequences and the x-axis is on a log2 scale. The data for all bacterial and archaeal phyla are available in Supplemental Tables 2 and 3, respectively.

### Focusing the census by environment

We were able to assign 89.3 and 95.1% of the sequences to one of seven broad environmental categories based on the metadata that accompanied the SILVA database (Tables 1). Across these broad categories there was wide variation in the number of sequences that have been sampled. Among bacterial sequences, the three best represented groups were from zoonotic (N=804,585), aquatic (N=214,085), and built environment (N=108,799) sources. Among the archaeal sequences the three best represented groups were the same, but ordered differently: aquatic (N=34,400), built environment (N=7,286), and zoonotic (N=5,597) (Figure 1C,D)). For both domains, soil samples were the fourth most represented category (bacteria: 74,870; archaea: 2,517). The orders of these categories was surprising considering soil and aquatic environments harbor the most microbial biomass and biodiversity (21). In spite of wide variation in sequencing depth and coverage (Table 1), the interquartile range across the fine-level categories for the bacterial OTU coverage only varied between 34.5 to 40.0 (median coverage=36.7%). The interquartile range in the OTU coverage by environment for the archaeal data was 41.5 to 53.1 (median coverage=44.9%). The archaeal coverage was higher than that of the bacterial OTU coverage for all categories except the food-associated, plant surface, and other invertebrate categories. Across all categories, the bacterial and archaeal sequencing data represented a limited number of phyla (Figure 4). Among the bacterial data, the fine-scale categories were dominated by Proteobacteria (N=24), Firmicutes (N=2), and Actinobacteria (N=1) and among the archaeal data, they were dominated by Euryarchaeota (N=16), Thaumarchaeota (N=10), and Aenigmarchaeota (N=1). Regardless, there were clear phylum-level signatures that differentiated the various categories. Within each of the bacterial and archaeal phyla, there was considerable variation in the relative abundance of each across the categories confirming that taxonomic signatures exist to differentiate different environments even at a broad taxonomic level.

**Figure 4.**
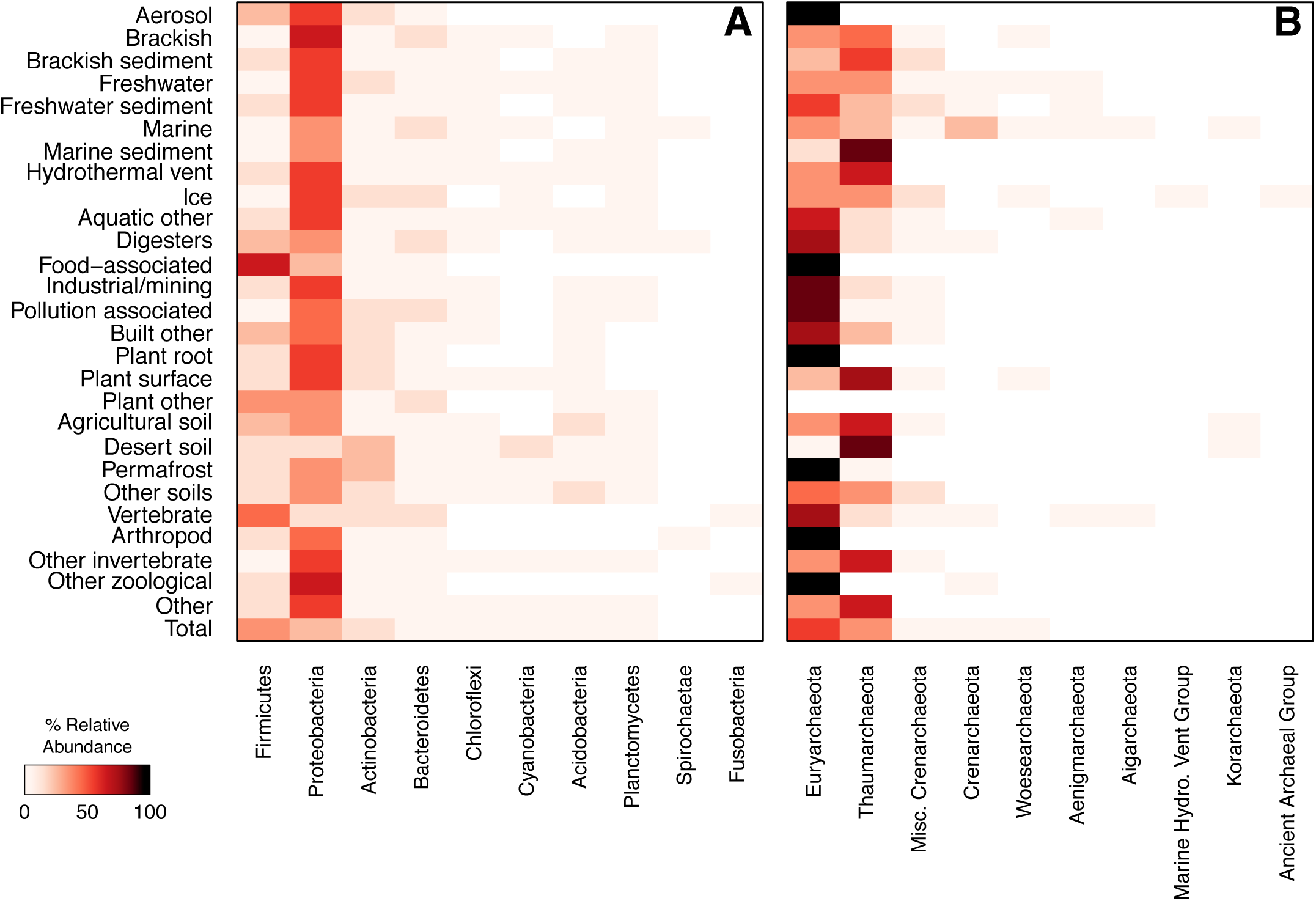
Heatmap depicting the relative abundance of the most common bacterial and archaeal phyla across different environments. Each environmental category exhibited a phylum-level signature although the bacterial census was dominated by sequences from the Firmicutes, Proteobacteria, Actinobacteria, and Bacteroidetes and the archaeal census was dominated by sequences from the Euryarchaeota and Thaumarchaeota. The ten most abundant phyla across all environmental categories are shown. The data for all bacterial and archaeal phyla are available in Supplemental Tables 4 and 5, respectively.

### The cultured census

In the 2004 bacterial census, there was great concern that although culture-independent methods were significantly enhancing our knowledge of microbial life, there were numerous bacterial phyla with no or only a few cultured representatives. To update this assessment, we identified those sequences that came from cultured and uncultured organisms. Overall, 18.9% of bacterial sequences and 6.8% of archaeal sequences have come from isolated organisms. Comparing the fraction of sequences deposited during and before 2006 from isolates to those collected after 2006, we found that culturing rates lag by 2.4 and 2.5-fold for bacteria and archaea, respectively. Among the 65 bacterial phyla, 24 have no cultured representatives and 14 of the 20 archaeal phyla have no cultured representatives. This lag is likely due to the differences in throughput of culture-dependent and-independent approaches. Of the phyla with at least one cultured representative, the median percentage of sequences coming from a culture was only 2.8% for the bacterial phyla and 1.7% for the archaeal phyla (Figure 5). Even though many phyla have cultured representatives, there is still a skew in the representation of most phyla found in cultivation efforts.

**Figure 5.**
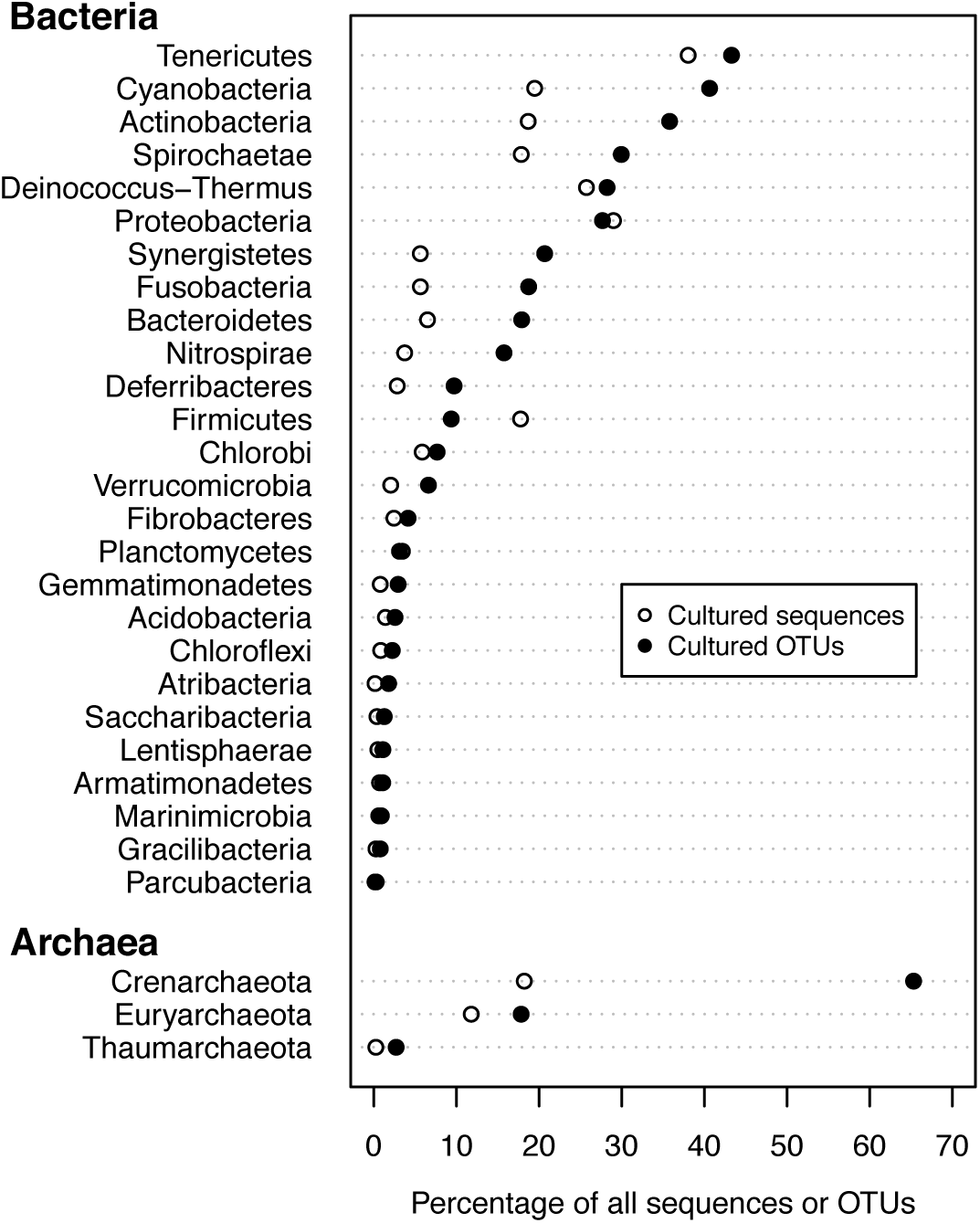
The rate that sequences and OTUs are generated from bacterial and archaeal cultures relative to all sequences and OTUs by phlum. Phyla with greater than 1,000 sequences are listed by domain. Open circles indicate the percentage of sequences in the database that match cultured organisms. Closed circles indicate the percentage of OTUs in this analysis that contain sequences belonging to a cultured organism. The data for all bacterial and archaeal phyla are available in Supplemental Tables 6 and 7, respectively.

Considering the possibility that large culture-independent sequencing efforts may only be re-sequencing organisms that already exist in culture, we asked what percentage of OTUs had at least one cultured representative. We found that 16.9% of the 117,385 bacterial OTUs and 13.1% of the 4,574 archeael OTUs had at least one cultured representative (Figure 5). Comparing the percentage of sequences with cultured representatives to the percentage of OTUs containing a sequence from a cultured representative revealed a strong cultivation bias within the Firmicutes, which had a higher percentage of sequences generated by cultivated representatives than would be expected based on the number of cultured organisms represented by OTUs (Figure 5). This likely reflects the extremely high number of cultivated biomedically relevant cultivars from genera such as *Bacillus, Streptococcus, Lactobacillus, Staphylococcus*, and others. Conversely, many phyla, including Cyanobacteria, Actinobacteria, Bacteroidetes, and Nitrospirae, had a lower percentage of sequences belonging to cultivated representatives than would be expected based on the percentage of OTUs that have sequences from cultured organisms, indicating that the cultivation efforts in these clades are relatively inefficient with regards to available diversity. Nevertheless, it is clear that the majority of OTUs from any phylum remain uncultivated, to say nothing of the diversity of organisms that may be encapsulated within the 97% sequence identity cutoff.

### New technologies to access novel biodiversity

Given the shift from Sanger sequencing to platforms that offer higher throughput but shorter reads, there is concern that our ability to harvest full-length sequences from communities will remain stalled. Several culture-independent methods have been developed that offer the ability to obtain full-length sequences of the 16S rRNA gene and even the complete genome. These have included single cell genomics (22) and assembly of short 16S rRNA gene fragments using data generated from PCR amplicons or metagenomic shotgun sequence data with the Expectation-Maximization Iterative Reconstruction of Genes from the Environment (EMIRGE) algorithm (23, 24). To test the ability of these technologies to expand our knowledge of microbial diversity beyond that of traditional approaches, we compared the overlap of OTUs found using each of the new methods with the traditional approaches (Figure 6). Utilizing the 16S rRNA gene sequences extracted from the single-cell genomes available on the Integrated Microbial Genomes (IMG) system (25), we identified 311 bacterial and 70 archaeal sequences, which were assigned to 115 and 27 bacterial and archaeal OTUs, respectively. Interestingly, only 8.7 and 3.7% of the bacterial and archaeal single celled OTUs, respectively, had not been observed by previous efforts. Next, we identified six studies that utilized EMIRGE to assemble 16S rRNA gene sequences from metagenomic sequences (23, 26–30). Together these studies assembled 599 bacterial and 9 archaeal full-length sequences, which were assigned to 335 and 7 bacterial and archaeal OTUs, respectively. Only 40.6 and 60.3% of the bacterial OTUs generated by this approach were previously identified by this traditional cultivation and PCR-based approaches, respectively. Although the application of this approach to Archaea has been limited, it was still surprising that 85.7 and 85.7% of the archaeal OTUs had been previously recovered by traditional cultivation and PCR-based approaches, respectively. Finally, we pooled 76,080 bacterial sequences from five studies that utilized EMIRGE to assemble 16S rRNA gene sequences from fragmented amplicons (24, 31–34). These sequences were assigned to 40,213 OTUs. We were surprised that only 7.6% of these OTUs were previously found by a more traditional approach. Although these PCR-based EMIRGE results may be valid, the high degree of novelty that was observed suggests that the error of the assembled reads may be too high for generating reference sequences. Each of these methods represent promising opportunities to continue the bacterial census using full-length sequences as well as genomic information.

**Figure 6.**
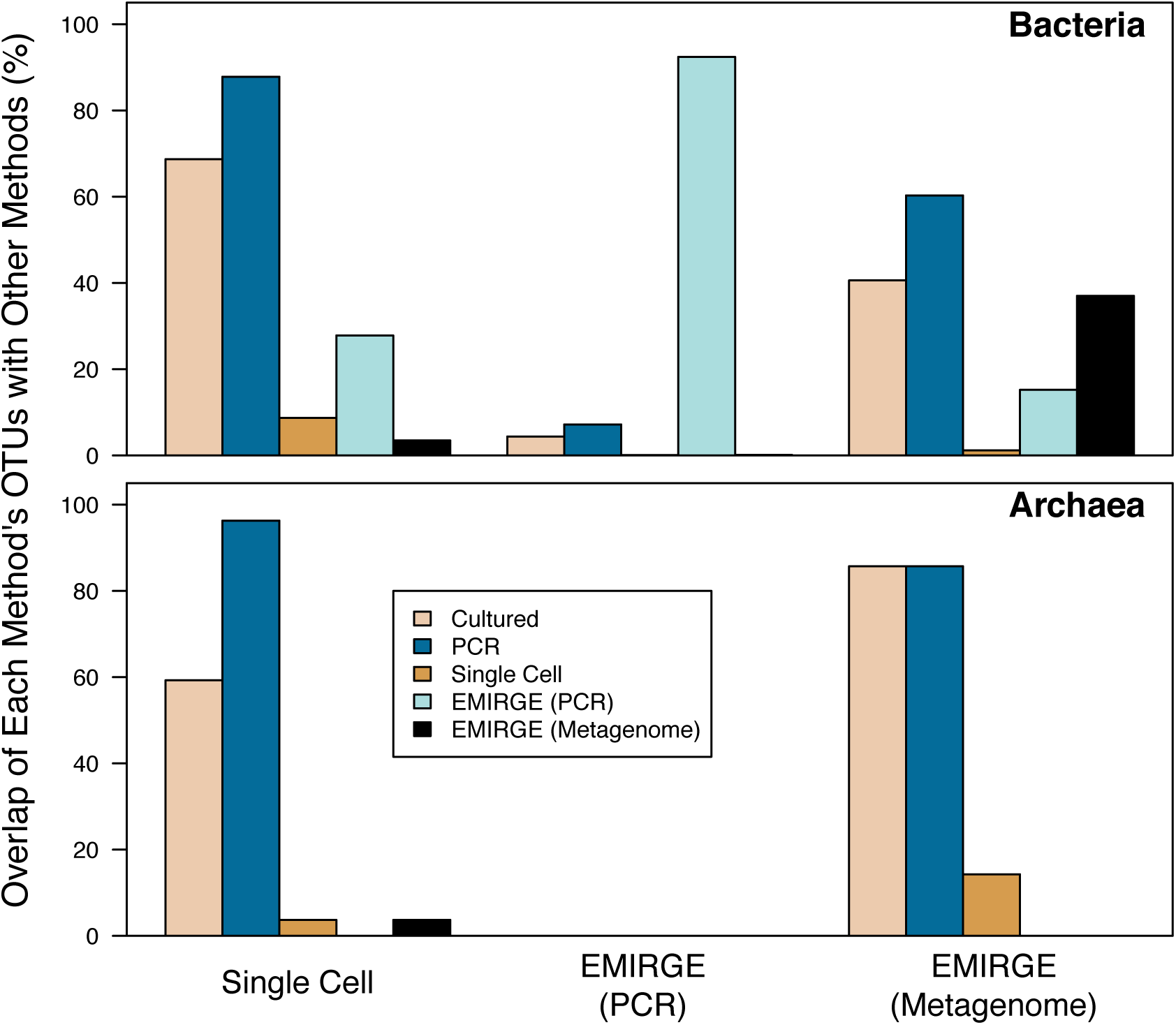
The percentage of bacterial and archaeal OTUs found by single cell genomics and EMIRGE using PCR or metagenomics that were also detected by other. The bars comparing a method to itself indicate the percentage of OTUs that were only detected by that method.

## Conclusions

It is clear that considerable biodiversity has been discovered since the first census in 2004. However, much of it has been biased towards particular phyla and environments. Our analysis suggests that 94.5% of new full-length bacterial and archaeal sequences are likely to have already been seen. Meanwhile, 29.2% of bacterial and 38.5% of archaeal OTUs have only been observed once. In spite of current estimates suggesting the global bacterial species richness may be as high as 10^12^ species (35), the current census based on full-length 16S rRNA gene sequences suggests that existing sampling methods will prevent us from acquiring full-length sequences for that level of diversity. As we have shown, current strategies repeatedly sample the same OTUs and do a poor job of resampling rarer populations. Given this low level of OTU coverage, it is likely that there are many more bacterial and archaeal populations yet to be sampled.

There are several additional reasons to suspect that the current census should be considered conservative. First, we found that most sequences recently deposited into public databases are being made by a small number of projects that have deeply sampled similar environments, and the number of full-length reads deposited into the databases has stalled. Second, it is widely acknowledged that 16S rRNA gene primers are biased; these biases are amplified when designing primers to amplify subregions used in sequencing short reads (36). Assembly of metagenomic data has shown the presence of introns in the 16S rRNA genes of organisms within the so-called “Candidate Phyla Radiation” (e.g. Saccharibacteria (TM7), Peregrinibacteria, Berkelbacteria (ACD58), WWE3 Microgenomates (OP11), Parcubacteria (OD1), et al.) that would preclude detection with standard PCR-based approaches (37, 38). Third, the willingness of researchers to contribute their sequences and the metadata describing the environment that the sequences were sampled from is critical for assessing the progress of the census and to accrue the benefits from having full-length sequences in the databases. Interestingly, the first 16S sequence was published in 1978, but was not available in a database until 1983. Similarly, only 5 of the 11 studies that used the EMIRGE algorithm deposited their sequences in GenBank. This makes the sequences from the other studies effectively invisible to the search algorithms used by 16S rRNA gene-specific databases to harvest sequences. As assembly and long read technologies advance, a mechanism is needed to assess the quality of the consensus sequences and to make them easily accessible to the 16S rRNA gene-specific databases.

Efforts to census microbial life using short read technology such as the International Census of Marine Microbes, the Earth Microbiome Project, and the Human Microbiome Project have significantly advanced our knowledge of microbial biogeography; however, these analyses have demonstrated the limitations of databases and taxonomies that are based on sequences from common and abundant organisms. During the period prior to the introduction of massively parallelized high throughput sequencing, it was common for a study to generate dozens or hundreds of sequences per sample. The existing databases that are used for classifying sequences are based on those sequences, which represent organisms that are generally abundant. We hypothesize that recent difficulties obtaining adequate classification for short sequences captured from more rare organisms are because our databases do not contain full-length references for those sequences. We fear that these trends will worsen unless researchers can leverage new sequencing and cultivation technologies to generate large numbers of full-length sequences from a large number of diverse samples.

Novel technologies such as single-cell genomics, metagenomics, and algorithms to recover full-length sequences from new sequencing platforms have demonstrated promise in circumventing previous limitations in identifying new OTUs. Using EMIRGE to assemble fragmented 16S rRNA gene amplicons may allow us to obtain deep coverage of communities; however, it is still unclear how faithful the assembled sequence is to that of the original organism. Additional sequencing technologies also offer the ability to directly generate full-length sequences, such as PacBio and potentially Oxford Nanopore. Initial application of PacBio to sequencing full-length fragments suggests that the sequences suffer from a high error rate (39). To obtain a more direct investigation of rare organisms, microbiologists are developing novel cultivation and single cell genomics techniques (???, 40–42). The ability to enrich or select for specific populations using these approaches could limit the need for redundant brute force sequencing. These approaches are still in active development, and we hope that through continuous refinement, they may allow us to significantly improve the coverage of OTUs in public databases.

## Materials and Methods

### Sequence data curation

The July 19, 2015 release of the ARB-formatted SILVA small subunit (SSU) reference database (SSU Ref v.123) was downloaded from http://www.arb-silva.de/fileadmin/silva_databases/release_123/ARB_files/SSURef_123_SILVA_19_07_15_opt.arb.tgz (43). This release is based on the EMBL-EBI/ENA Release 123, which was released in March 2015. The SILVA curators identify potential SSU sequences using keyword searches and sequence-based search using RNAmmer (http://www.arb-silva.de/documentation/release-123/). The SILVA curators then screened the 7,168,241 resulting sequences based on a minimum length criteria (<300 nt), number of ambiguous base calls (>2%), length of sequence homopolymers (>2%), presence of vector contamination (>2%), low alignment quality value (<75), and likelihood of being chimeric (Pintail value < 50). Of the remaining sequences, the bacterial reference set retained those bacterial sequences longer than 1,200 nucleotides and the archaeal reference set retained those archaeal sequences longer than 900 nucleotides. The aligned 1,515,024 bacterial and 59,240 archaeal sequences were exported from the database using ARB along with the complete set of metadata. Additional sequence data was included from single-cell genomes available on the Integrated Microbial Genomes (IMG) system (25), many of which were recently obtained via the GEBA-MDM effort in Rinke et al. (22). “SCGC” was searched on the IMG database March 12, 2015 to download the bacterial (N=249) and archaeal (N=46) 16S rRNA gene sequences and their associated metadata. Further, sequences generated from amplicon and shotgun metagenomic data using the EMIRGE program were also included (23, 24). The IMG and EMIRGE sequences were aligned against the respective SILVA-based reference using mothur (44). The aligned bacterial and archaeal sequence sets were pooled and processed in parallel. Using mothur, sequences were further screened to remove any sequence with more than 2 ambiguous base calls and trimmed to overlap the same alignment coordinates. The sequences in the resulting bacterial dataset overlapped bases 113 through 1350 of an *E. coli* reference sequence (V00348) and had a median length of 1,233 nt. The sequences in the resulting archaeal dataset overlapped positions 362 to 937 of a *Sulfolobus solfataricus* reference sequence (X03235) and had a median length of 580 nt. The archaeal sequences were considerably shorter than their initial length because it was necessary to find a common overlapping region across the sequences. The final datasets contained 1,411,234 bacterial and 53,546 archaeal 16S rRNA gene sequences. Sequences were assigned to OTUs using the average neighbor clustering algorithm (45).

### Metadata curation

The metadata that was contained within the SSU Ref database was used to expand our analysis beyond a basic count of sequences and the number of OTUs in each domain. The environmental origins of the 16S rRNA gene sequences were manually classified using seven broad “coarse” categories, and further refined to facilitate additional analyses with twenty-six more specific “fine” categories (Table S1). These were assigned based on manual curation of the “isolation_source” category within the ARB database associated with each of the sequences. For source definitions that were not identifiable by online searches, educated guesses were made or they were placed into the coarse “Other” category. There were 151,669 bacterial and 2,565 archaeal sequences where an “isolation_source” term was not collected. We ascertained whether a sequence came from a cultured organism by including those sequences that had data in their “strain” or “isolate” fields within the database and excluded any sequences that had “Unc” as part of their database name as this is a convention in the database that represents sequences from uncultured organisms. Complete tables containing the ARB-provided metadata, taxonomic information, OTU assignment, and our environmental categorizations are available at FigShare for the bacterial (https://dx.doi.org/10.6084/m9.figshare.2064927) and archaeal (https://dx.doi.org/10.6084/m9.figshare.2064942) data.

### Calculating coverage

Sequencing coverage (C_Sequence_) was quantified by two methods. The first was to use Good’s coverage according to

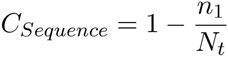

where n_1_ is the number of OTUs represented by only one sequence and N_t_ is the total number of sequences (46). Although Good’s coverage provides information about the success of the sequencing effort in sampling the most abundant organisms in a community, it does not directly provide information about the success of the sequencing effort in recovering previously unobserved OTUs. To quantify the ability of sequencing to identifying novel OTUs or, in other words, to quantify the “distance” in the peak of the rarefaction curves to their hypothetical asymptote, we defined “OTU coverage” (C_OTU_) as

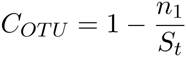

where S_t_ is the total number of OTUs. Whereas Good’s coverage estimates the probability that a new sequence will have already been seen, OTU coverage estimates the probability that a new OTU will match an existing one. It is therefore an extension of Good’s coverage in that it quantifies the probability that, for any given set of sequences clustered into an OTU, that OTU will have already been seen. Thus, high Good’s coverage means that any new sequence is unlikely to be novel, and high OTU coverage means that any new OTU is unlikely to be novel.

### Data analysis

Our analysis made use of ARB (OS X v.6.0) (43), mothur (v.1.37.0) (44), and R (v.3.2.2) (47). Within R we utilized the knitr (v.1.10.5), wesanderson (v.0.3.3.99), and openxlsx (v. 2.4.0) packages. A reproducible version of this manuscript including data extraction and processing is available at https://www.github.com/SchlossLab/Schloss_Census2_mBio_2016.

